# The dependency of TMS-evoked potentials on electric-field orientation in the cortex

**DOI:** 10.1101/2025.01.10.632309

**Authors:** Ida Granö, Tuomas P. Mutanen, Aino E. Nieminen, Jaakko O. Nieminen, Victor H. Souza, Risto J. Ilmoniemi, Pantelis Lioumis

## Abstract

Transcranial magnetic stimulation (TMS) stimulates the brain by electromagnetic induction. The outcome depends on multiple stimulation parameters such as the induced electric-field pattern (in particular, the location of the peak field and its orientation), intensity and timing. However, it is not clear how the TMS-evoked responses are affected by all the stimulation parameters. This study elucidates the dependency of the TMS-evoked electroencephalography (EEG) responses on the orientation of the stimulating electric field. To achieve this, we analysed a dataset from six subjects who were given pulses with 36 stimulus orientations to the pre-supplementary motor area (pre-SMA). The TMS-evoked potentials (TEPs) and induced oscillations were analysed with cluster-based statistics. Source estimation was performed to assess the effects of stimulus orientation on the TMS-evoked signal propagation. The amplitudes of the early peaks (20 and 40 ms after the TMS pulse) strongly depended on the electric field orientation. Our analysis suggested orientation dependency up to 100 ms post-stimulus in most subjects, indicating changes in stimulation efficacy and potential changes in signal propagation from the stimulated site. These results suggest that different orientations may perturb different networks. Thus, the orientation is a crucial parameter for the stimulation outcome and should be adjusted according to the cortical network under investigation.

## 1 Introduction

Transcranial magnetic stimulation (TMS) induces an electric field inside the brain, triggering neuronal action potentials. However, applying TMS is not always straightforward due to the many stimulation parameters, such as the location and orientation of the peak induced electric field, the overall stimulation intensity, and the timing of the TMS pulse. All these parameters significantly affect the stimulation outcomes [1,2], yet their roles in shaping the TMS-evoked responses are not yet well known. Various studies have aimed to shed light on this issue through the systematic recording of the physiological responses resulting from various stimulation parameters [3–10] and through computer simulations [11–16]. Understanding the effects of different stimulation parameters helps target specific cortical circuits, increasing the utility of TMS as a functional and structural neuroanatomical tool for mapping distinct cortical regions.

The stimulus orientation is generally thought to influence the efficacy of TMS [13,15,17] and might affect which neurons are primarily stimulated by the TMS pulse [11,16,18,19]. As TMS-evoked EEG responses are sensitive to even slight coil shifts and rotations [1,20], precise work has to be done to investigate the effects of the stimulus orientation on TMS-evoked potentials (TEPs).

The aim of this work was to determine how the amplitudes, frequency-spectral properties and effective connectivity patterns of TEPs in time and space depend on stimulus orientation. For this purpose, we analysed a dataset [20] containing EEG responses to TMS delivered in 36 different orientations in six subjects. We expected that at least the early (< 50 ms after the TMS pulse) TEP amplitudes at the stimulation site would be affected by changes in stimulus orientation, in line with earlier work [20] that focused on these early responses. However, stimulus orientation could also be reflected in overall activation patterns at later times also further from the stimulated site. It has been shown that the orientation affects TMS-induced functional connectivity [8], indicating that the orientation plays a role also at longer time scales. This work explores the effect of stimulus orientation on the spatiotemporal domain of the TEPs and on effective connectivity elucidated by TMS–EEG.

## 3 Methods

### 3.1 Data acquisition and pre-processing

The data used in this work were originally obtained in [20] and acquired in the following way:

Six subjects (referred to as S1–S6) participated in the study (2 males, one left-handed, ages 22– 42 years). Before the experimental session, T1-weighted, fat-suppressed T1-weighted, and T2-weighted MRI images were recorded of each subject.

The brain activity was recorded with a BrainAmp DC amplifier (Brain Products GmbH, Germany), with a low-pass filter with a 1000-Hz cut-off frequency, sampled at 5000 Hz. Subjects wore earbuds playing white noise mixed with recorded TMS coil click sounds [21] underneath headphones to suppress the perception of the TMS coil click. The sound levels were adjusted to maximally mask the click sound while being within the safety and comfort limits of the subject.

An mTMS system [22] with a 2-coil set [9] capable of electronically adjusting the stimulus orientation was used to deliver trapezoidal monophasic pulses to the left pre-supplementary motor area (pre-SMA; [23]). This allowed control of stimulus orientation with 1-degree precision without moving the coil physically, removing variations in coil tilt and reducing repositioning errors. A thin foam was attached to the bottom of the coil to reduce bone conduction of the TMS coil click sound. An eXimia NBS 3 neuronavigation system (Nexstim Plc, Finland) ensured targeted and consistent stimulation throughout the experimental session. The coil was placed over the superior frontal gyrus, slightly anterior to the vertical anterior commissure [23]. The stimulation intensity was adjusted to produce a peak-to-peak amplitude of 5–10 µV in the 15–50 ms complex [24]. Forty-eight pulses were given in evenly spaced 36 orientations in 12 randomized blocks, with a pseudo-randomized interstimulus interval sampled from a uniform distribution between 2.4 and 2.7 s, resulting in 1728 pulses per subject. The 0° orientation was defined as being parallel to the stimulated gyrus.

Data were pre-processed in Matlab R2020 or newer (MathWorks Inc., USA). First, the TMS pulse artefact (–2…8 ms) was cut from each trial (–600…600 ms) and replaced with values obtained with cubic interpolation. A high-pass zero-phase-shift Butterworth filter with a 1-Hz cut-off frequency was applied. Then, trials heavily contaminated by eye blinks or scalp muscle activation were removed from further analysis. The data were baseline corrected by subtracting from each signal the average signal value in the same channel in the time range –200…–10 ms. The SOUND algorithm [25] was applied to remove noise, utilizing individual head geometry. Finally, the signal in each channel was low-pass filtered with a 45-Hz cut-off frequency, resampled to 1000 Hz, and expressed as the voltage with respect to the average potential among all channels at each moment of time.

Before pre-processing, S1 showed large muscle artefacts right after the TMS pulse, while S3 had persistent muscle activity. S4 had line noise and some stimulation artefacts in the data, while S6 had large decay artefacts in channel FCz. On visual inspection, the preprocessing effectively removed these disturbances from the data.

### 3.2 Data analysis

#### 3.2.1 TEP analysis

To explore the effects of the stimulus orientation on the TEP amplitudes in the spatiotemporal domain, we applied a cluster-based statistical approach to the data with the Fieldtrip toolbox [26]. The analyses were run on both an intra-subject and inter-subject level. First, the datasets from each subject were analysed with cluster-based statistics. We used single-factor ANOVA as the initial test statistic to test the effect of the stimulus orientation on TEP amplitudes at each channel-time point. To increase the power of the initial statistic test, neighbouring orientations were paired into single distributions to increase within-group sample size, and ANOVA was then run on the resulting 18 orientations instead of the original 36, with 75–96 (mean 89, sd 4.5) observations at each grouped orientation. We tested three different strategies to merge the subject-level statistics into group-level statistics: Fisher’s method [27], Stouffer’s Z method [28], and combining the data from all subjects into one large dataset [29]. For details on the application of Fisher’s and Stouffer’s Z methods, see Appendix A.

Before running the tests, the data were checked against the test assumptions (normality, homoscedasticity, independence). Normality was assessed by normal probability plots, and variances across orientations were examined visually by boxplots (Supplementary material). The trials were assumed to be independent.

For all tests run with cluster-based statistics, the threshold for clustering was set at *p* = 0.05, and a time range of 0–500 ms was examined. The tests were run with 1000 randomizations. For the individual tests, the cluster-level alpha value was set at (1 – *r* × 0.05/6) according to the Benjamini–Hochberg procedure [30], where *r* is the rank of the dataset and 6 is the number of subjects. For the meta-analyses, the cluster-level alpha was set at 0.05.

#### 3.2.2 Time–frequency analysis

To assess differences in the time–frequency spectra across different stimulus orientations, the event-related spectral perturbation (ERSP; [31]) was computed for each trial with a multitaper time–frequency transformation. A 400-ms-long Hanning window was utilized as the taper, and the ERSP was computed from 2 to 45 Hz in the time window –500…500 ms around the TMS pulse with –500…–200 ms as baseline.

As in the TEP analysis, cluster-based statistics were computed for each subject with ANOVA as the initial test statistic. Statistics were computed both individually for each subject, and by combining subjects with Fisher’s method, Stouffer’s Z method, and a combined dataset for all subjects. The cluster-level alpha values were set as in the TEP analysis.

#### 3.2.3 Source analysis

Source estimation was performed to investigate further changes in the propagation of TMS-evoked activity after giving pulses with different electric-field orientations. Each depth-weighted minimum-norm-estimated (MNE; [32]) source point was thresholded against a permuted baseline threshold. The threshold was calculated by permuting the baseline across the conditions 1000 times by drawing random trials equal to the condition with the least trials remaining after preprocessing (range: 75–91 trials). Then, the maximum over the whole spatiotemporal domain of the MNE estimate of each permutation was stored. The threshold was set at 95^th^ percentile of the stored maxima.

To investigate changes in the source estimates when stimulating in different orientations, we evaluated the Activation Concurrence, i.e., the number of conditions that produced suprathreshold sources in any given spatiotemporal point, as well as the Activation Variability, i.e., the variance over the conditions, normalized by the maximum variance per subject. The Activation Concurrence and Activation Variability were then visualized in 10-ms intervals for qualitative analysis of differences in signal spread between the conditions.

Subject-specific lead fields were calculated by extracting the scalp, skull and white matter from individual MRIs with the SimNIBS software [33] and utilizing the boundary element method for calculating the lead-field matrices. The regularization parameter for the MNE estimates were set for each individual with Morozov’s discrepancy principle. The found regularization parameters ranged from 0.08 to 0.3. The minimum-norm estimates were depth-weighted [34] with a depth parameter of 0.3, and the dipoles were placed with a free orientation in the lead field matrix. See Appendix A for more details.

## 3 Results

### 4.1 Visual inspection

Visual inspection suggested orientation dependency in the amplitude of the 20- and 40-ms peaks in all subjects (Figure 1). This dependency was most evident close to the stimulated site, and roughly followed a sinusoid with a period of 180° and the largest amplitudes at –90° and 90°. Two subjects (S3 and S6) also showed some orientation dependency in the 20- and 45-ms peaks over the primary motor cortex, as seen in electrode C3. Most subjects also displayed an orientation dependency in the 6-ms peak, with a larger negative deflection at 90°. Later peaks showed no clear changes in amplitudes with different stimulus orientations.

**Figure 1.**
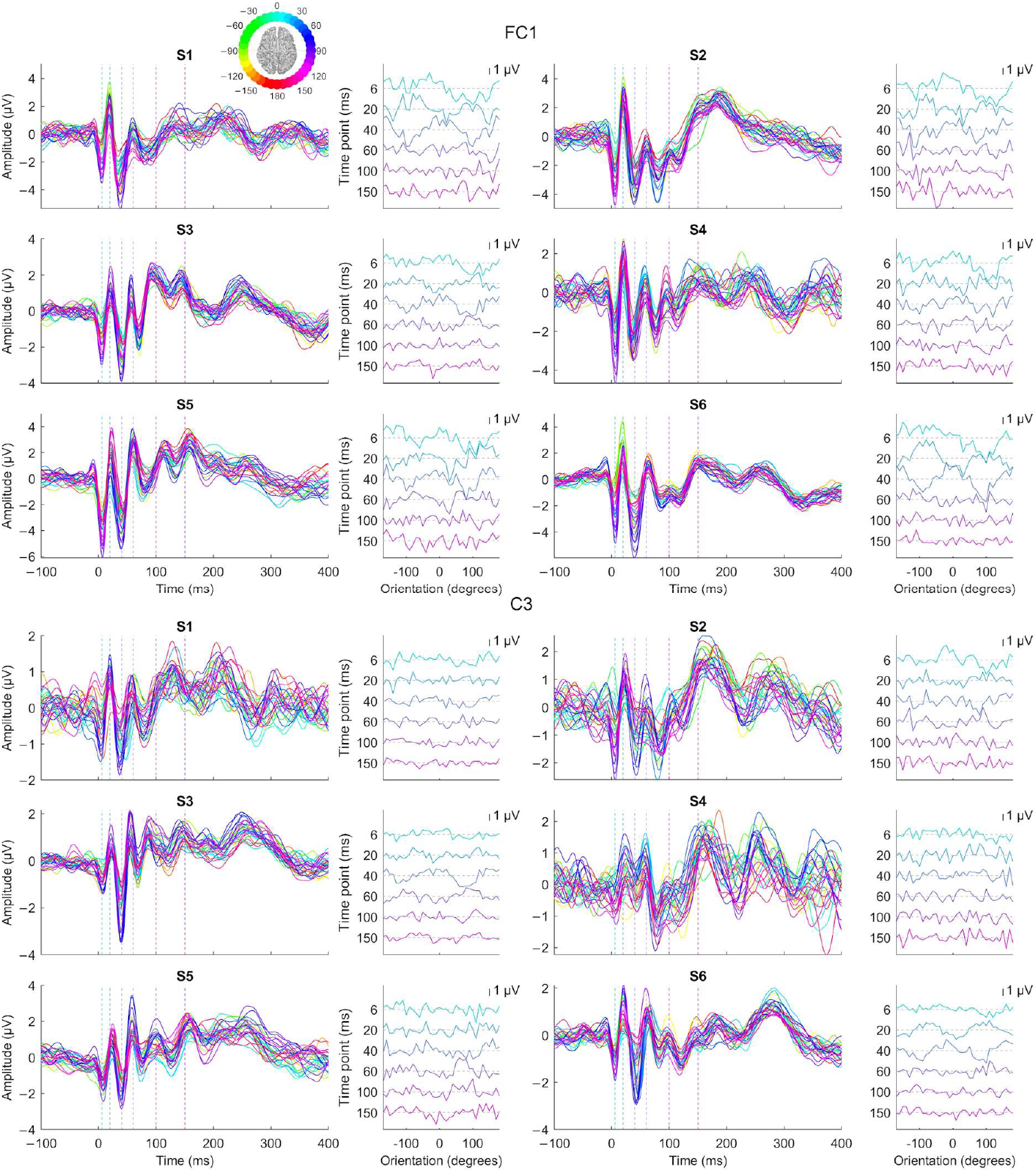
TMS-evoked potentials (TEP) close to the stimulation site (channel FC1) and over the left primary motor cortex (C3) for all six subjects. On the right of each TEP plot is the mean amplitude of each orientation plotted against the angle at chosen time points (6, 20, 40, 60 100 and 150 ms post-stimulus). The compass in the upper left corner shows the orientation for each TEP by colour.

The visual inspection of topographies as a function of time and orientation implied that the voltage patterns of the neuronal responses to TMS stayed consistent across the different stimulus orientations (see Supplementary material). The topographies that showed clear difference to the overall patterns were associated with low amplitude.

### 4.3 Cluster-based statistics on TEPs

Five out of six subjects (S1, S2, S3, S5 and S6) showed a significant effect of stimulus orientation on TEP amplitudes (Figure 2), while S4 showed a strong trend (*p* = 0.052), according to the cluster-based statistical analysis. In four subjects, the clusters ended before 100 ms and were mainly focused at and close to the stimulation site. In S3, the cluster extended to 184 ms. In S2, S3 and S6, differences were also found around the ipsilateral motor cortex (C3). S2 and S3 also displayed differences across stimulus orientations in channels over the contralateral hemisphere. The time ranges and *p*-values of the significant clusters are shown in Table 1.

**Table 1.**
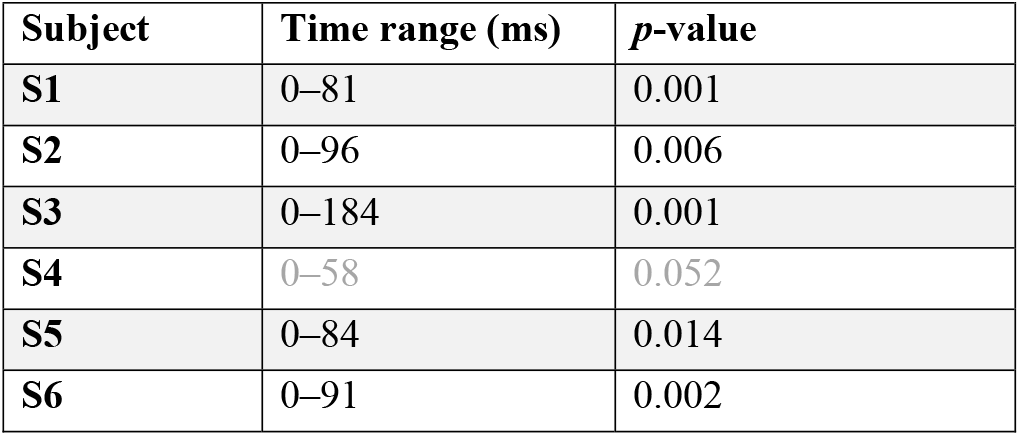
The time ranges and *p*-values for the significant clusters for each subject, as well as the largest clusters for S4.

**Figure 2.**
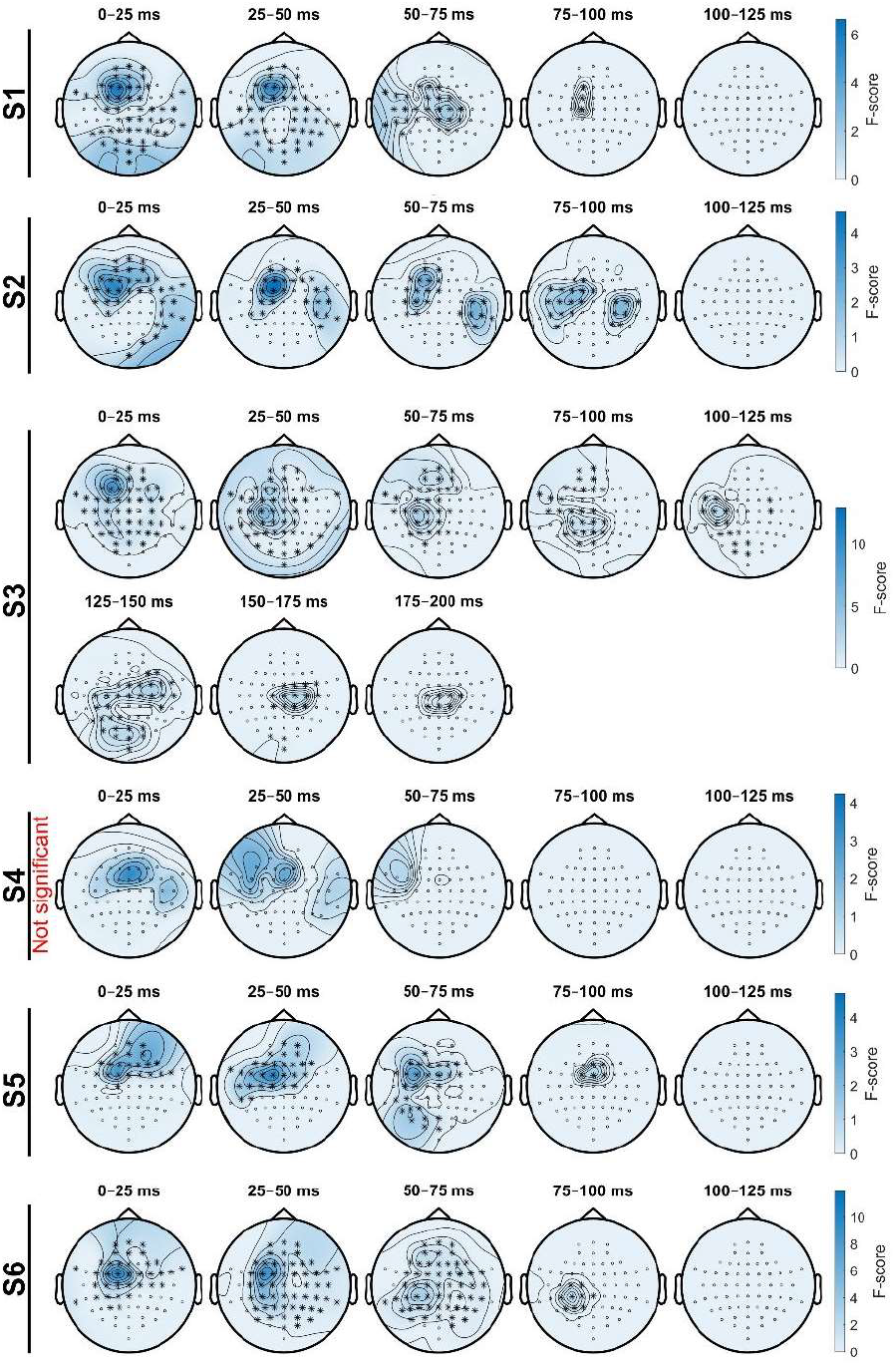
The cluster-based statistics suggest that the stimulus orientation affects TEP amplitudes until approximately 100 ms in most subjects. The significant clusters are marked by asterisks when ANOVA was used as the initial test statistic. Subject S4 showed no significant changes in TEPs across orientations. The darker blue areas show the values of the test statistic within the largest cluster for each subject, indicating the relative strengths of the effects.

Combining the subject data using Fisher’s method and Stouffer’s Z method, and by combining all datasets together gave comparable results, as shown in Figure 3. In all three cases, the clusters were centred around the stimulation site, and at early latencies (<25 ms), the clusters suggested changes in channels across the whole head. Fisher’s method resulted in the largest cluster, indicating changes in TEPs up until 193 ms. Table 2 shows the *p*-values and ranges of the significant clusters.

**Table 2.** The *p*-values and time ranges of the significant clusters when using Fisher’s method, Stouffer’s Z method, and combining all datasets together to assess the inter-subject effects of orientation on TEPs.

| Method | Time range (ms) | $p$-values |
| --- | --- | --- |
| **Fisher's method** | 0–193 | 0.001 |
| **Stouffer's Z method** | 0–114 | 0.001 |
| **Combining datasets** | 0–65 | 0.001 |

**Figure 3.**
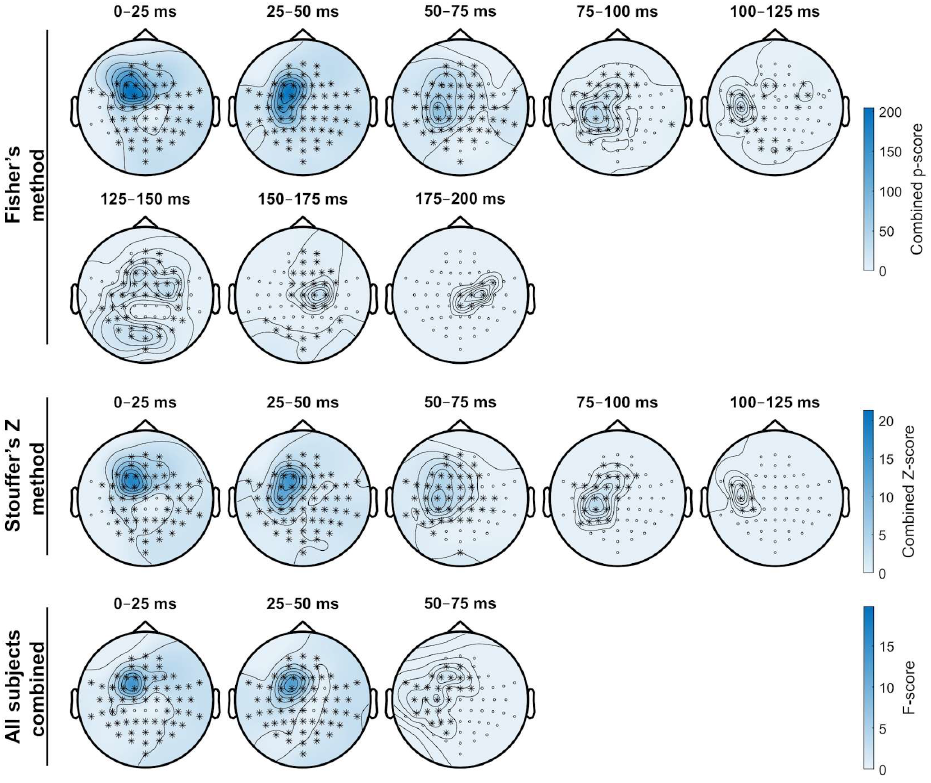
When combining the data from all subjects, the results were similar to those from the individual subjects. The clusters are marked by asterisks, when the data from the subjects were combined with Fisher’s method, Stouffer’s Z method, and by combining the data from all subjects into one big dataset. The dark blue areas show the test statistic values within the clusters, indicating the relative strengths of the effects.

Visually inspecting the TEPs in the electrodes with the largest test statistic in 25 ms blocks indicate that, at early time points (< 15 ms), a stimulus orientation of approximately 100° produced a larger negative deflection at the stimulated site in most subjects, after which stimulus orientations around –90 and 90° produced the strongest responses (Figure 4). The largest differences across orientations did not remain at the stimulated site, but spread to other nearby areas, such as towards the motor cortex, to parietal areas, and to the contralateral hemisphere. The different approaches to combining the datasets resulted in small differences in the exact location of the largest differences across the orientations.

**Figure 4.**
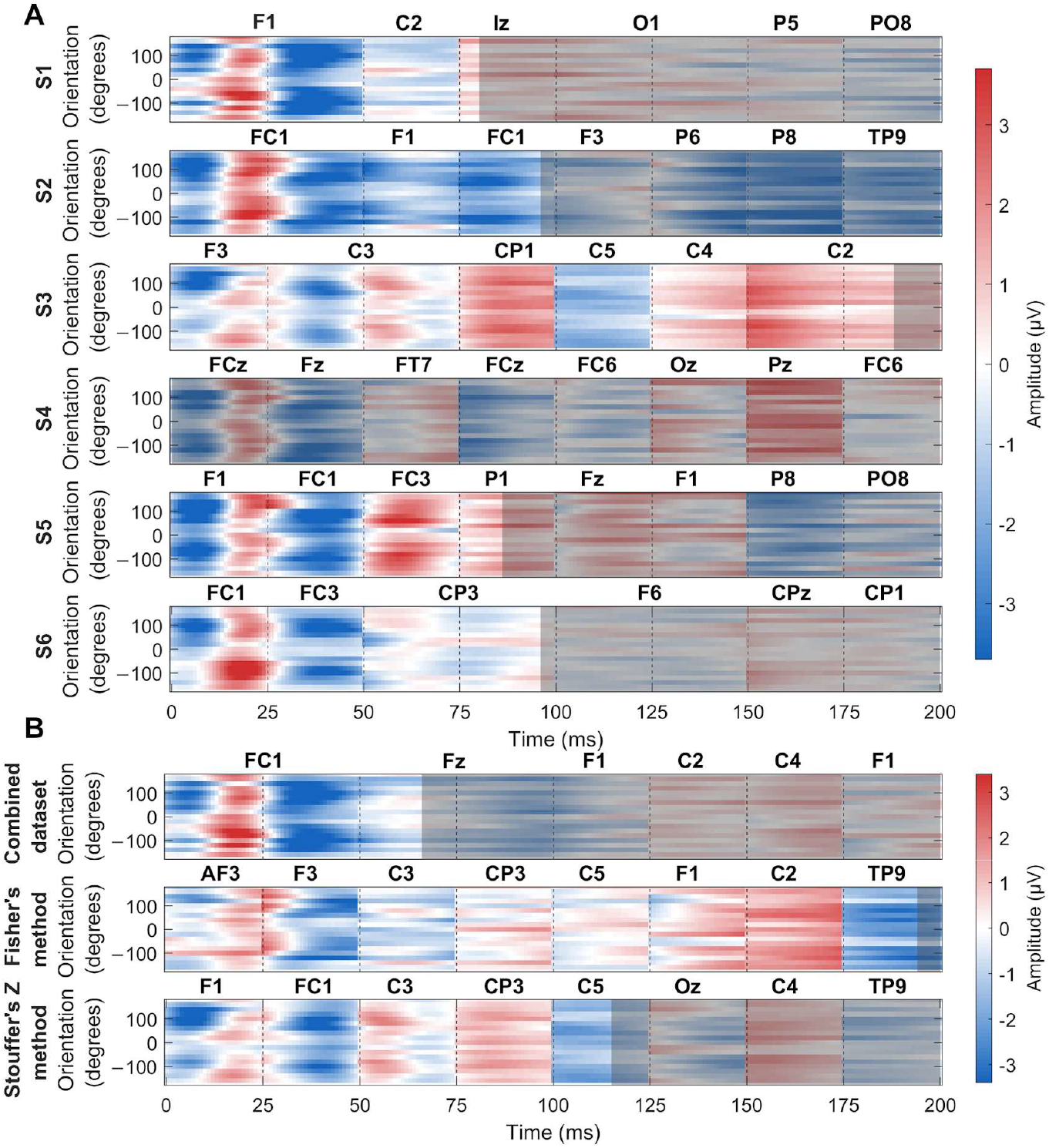
Visually inspecting the data over the orientations in the channel with the largest test statistic in 25-ms time windows suggests that early responses (< 15 ms) show largest negative peaks close to the stimulated site at around 100° in many subjects, after which orientations centred around –90 and 90° appear to evoke the strongest responses. After 50 ms, the statistical analysis suggests that the largest differences across orientations are found further from the stimulated site, such as centrally or parietally. Time windows outside of significant clusters are masked with grey.

#### Time–frequency analysis

All six subjects showed a significant effect of stimulus orientation (Figure 5) on the ERSP according to the cluster-based statistical analysis. In all subjects, the largest differences in ERSP were in the beta (13–30 Hz) and gamma bands (30–70 Hz) close to the stimulation site (F1, FC3, FC1). In S3, S5 and S6, the changes extend to C1, C3, CP1 and CP3. In S4, the largest effect was in FCz. The cluster-based analysis suggested changes in ERSP starting at 5 Hz in S6, at early beta in S1, S2 and S3, and in mid-range beta in S4 and S5. In all subjects, most of the clusters were contained within the first 150 ms after stimulation. The exact frequency ranges and *p*-values of the significant clusters are shown in Table 3.

**Table 3.**
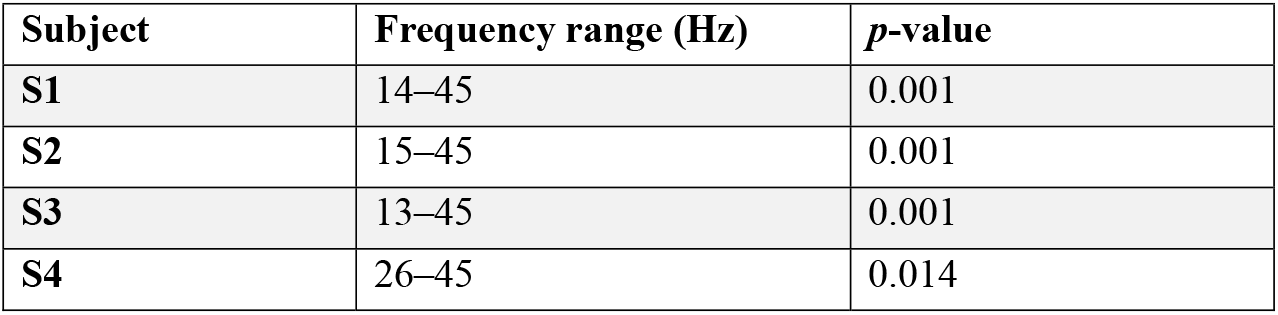
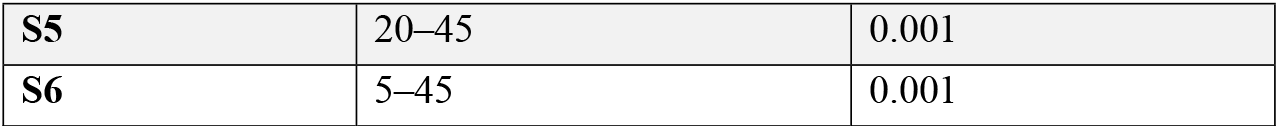
The time ranges and *p*-values for the significant clusters for each subject.

| Subject | Frequency range (Hz) | $p$-value |
| --- | --- | --- |
| S1 | 14–45 | 0.001 |
| S2 | 15–45 | 0.001 |
| S3 | 13–45 | 0.001 |
| S4 | 26–45 | 0.014 |
| **S5** | 20–45 | 0.001 |
| **S6** | 5–45 | 0.001 |

**Figure 5.**
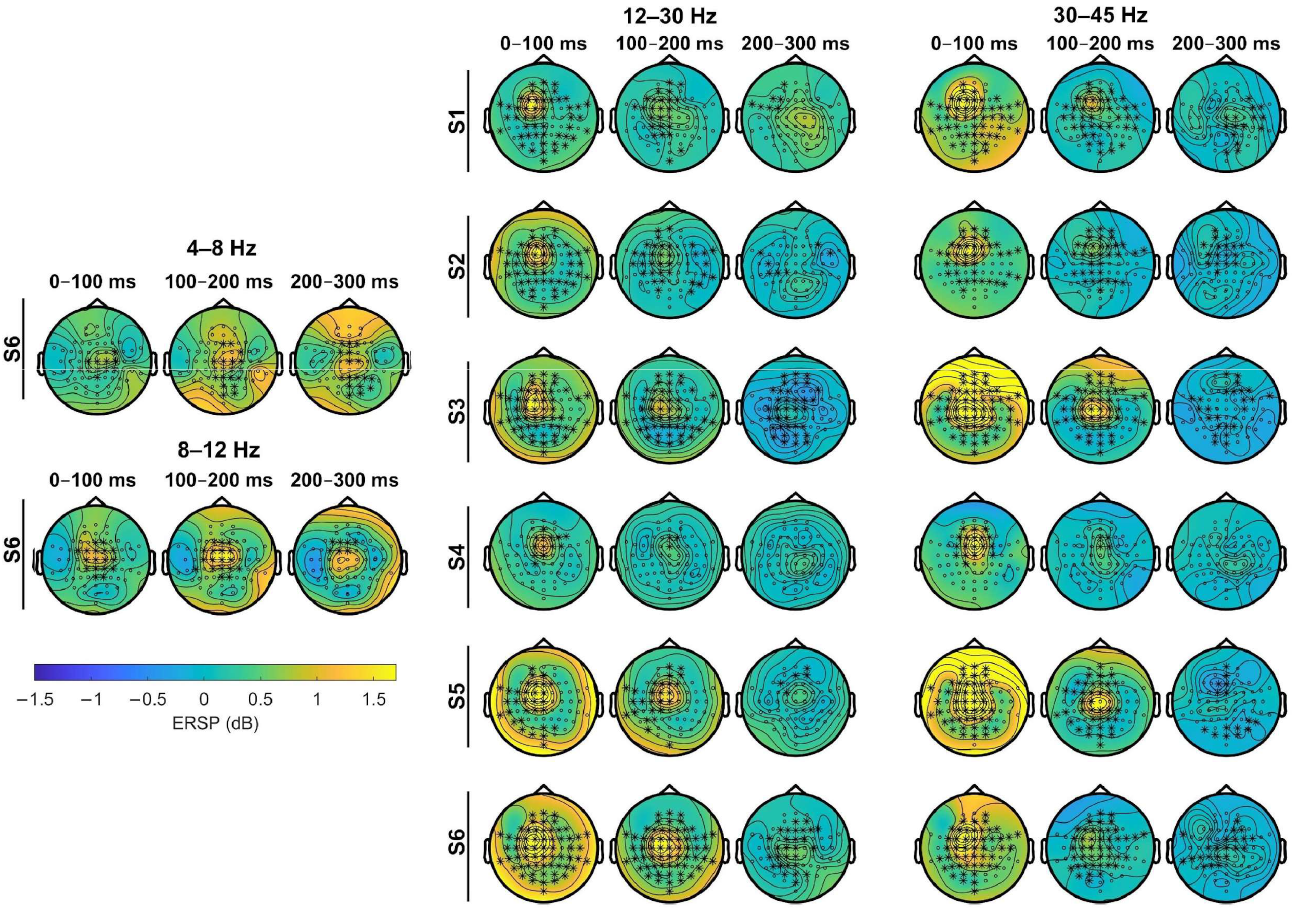
The stimulus orientation affected the induced activity in the beta-gamma range in all subjects. The significant clusters are marked by asterisks for each subject separately. The colour indicates the bandpower of the signals with respect to baseline (–500…–200 ms) in the specific frequency bands (4–8 Hz, 8–12 Hz, 12–30 Hz, and 30–45 Hz) averaged across 100 ms time windows.

Combining the subject data using Fisher’s method and Stouffer’s Z method, and by combining all datasets together gave quite similar results, as shown in Figure 6, although with small differences in the exact extent of the clusters. Again, Fisher’s method resulted in the largest cluster, spanning the whole computed frequency range (1–45 Hz). In all three cases, the clusters were centred around the stimulation site. Table 4 shows the *p*-values and frequency ranges of the significant clusters.

**Table 4.** The *p*-values and frequency ranges of the significant clusters when using Fisher’s method, Stouffer’s Z method, and combining all datasets together to assess the inter-subject effects of orientation on ERSP.

| Method | Frequency range (Hz) | *p*-values |
| --- | --- | --- |
| Fisher's method | 1–45 | 0.001 |
| Stouffer's Z method | 13–45 | 0.001 |
| Combining datasets | 12–45 | 0.001 |

**Figure 6.**
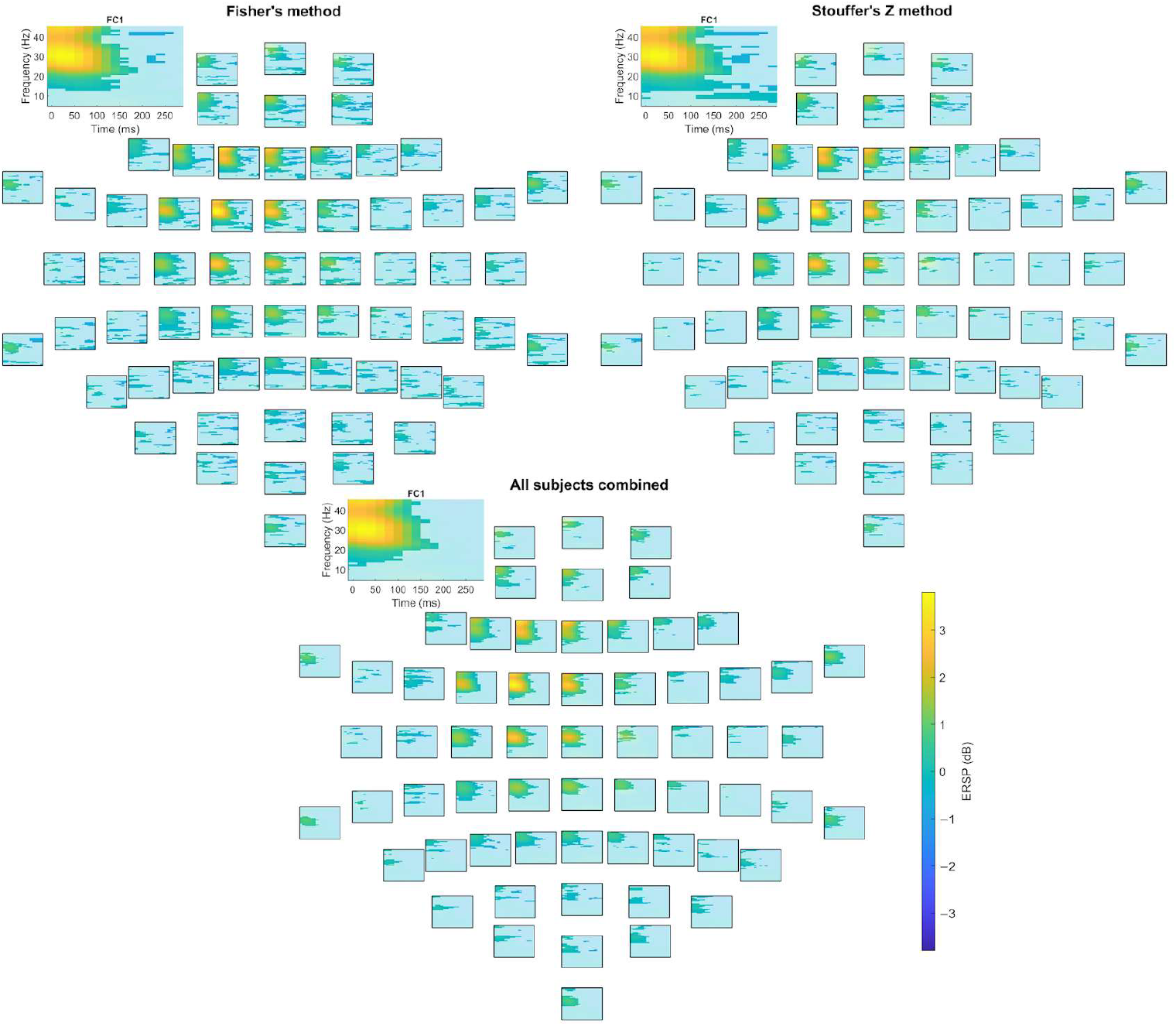
The results of the cluster-based statistical test for the subject-combined statistics were comparable to those run on the individual subjects. The colour indicates the bandpower of the signals from all the subjects with respect to baseline (–500…–200 ms). Areas outside of the significant clusters are masked.

#### Source estimates

The source estimates suggested some orientation-dependent differences in signal propagation from the stimulated site. Figure 7 shows the Activation Concurrence, i.e., the number of suprathreshold sources over the different orientations, as well as the Activation Variability, i.e., the normalized standard deviation across orientations at different source and time points. Early peaks (5–15 ms) were most prominent in S2 and S5. The results were highly consistent across the subjects at 35–45 ms after the TMS pulse, showing both high Activation Concurrence and Activation Variability, indicating that the response is stable, yet the amplitude varies with orientation. After 45 ms, the heterogeneity in signal propagation increased: S2, S3, S4 and S6 had later suprathreshold responses in some conditions centrally, S5 showed suprathreshold responses over M1 at approximately 55–65 ms, and S1 had small suprathreshold responses past 55 ms.

**Figure 7.**
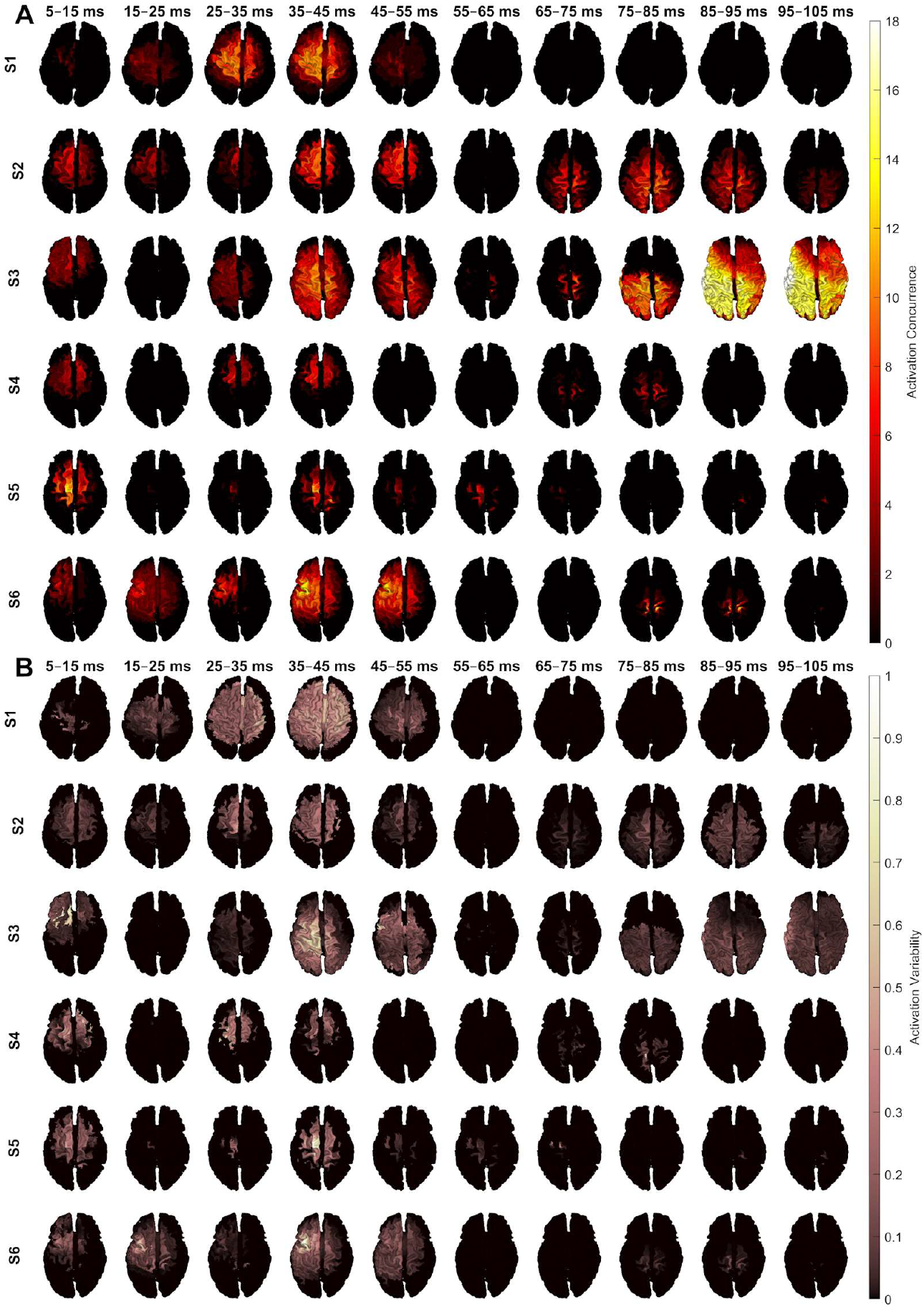
We estimated the effect of the stimulus orientation on effective connectivity. We quantified both the Activation Concurrence (A), i.e., how often different E-field orientations result in suprathreshold cortical activation, and the Activation Variability (B), i.e., how much the observed suprathreshold activations vary in amplitude across the orientation. On the colormap, black indicates no suprathreshold sources, relating to low Activation Concurrence. At 35–45 ms, both the Activation Concurrence and Activation Variability are high in most subjects at the stimulated site, after which the heterogeneity in signal propagation from the stimulated site increases.

The orientations producing suprathreshold sources are shown in Figure 8 from the source point with maximum Activation Concurrence at each time point. In the 5–15-ms time window, orientations around 100° were most likely to produce suprathreshold sources. In the 15–50-ms time window, on the other hand, orientations near –90 and 90° produced suprathreshold sources most consistently. After 50 ms, inter-subject variability increased, resulting in larger spreads of orientations producing suprathreshold sources at different time intervals.

**Figure 8.**
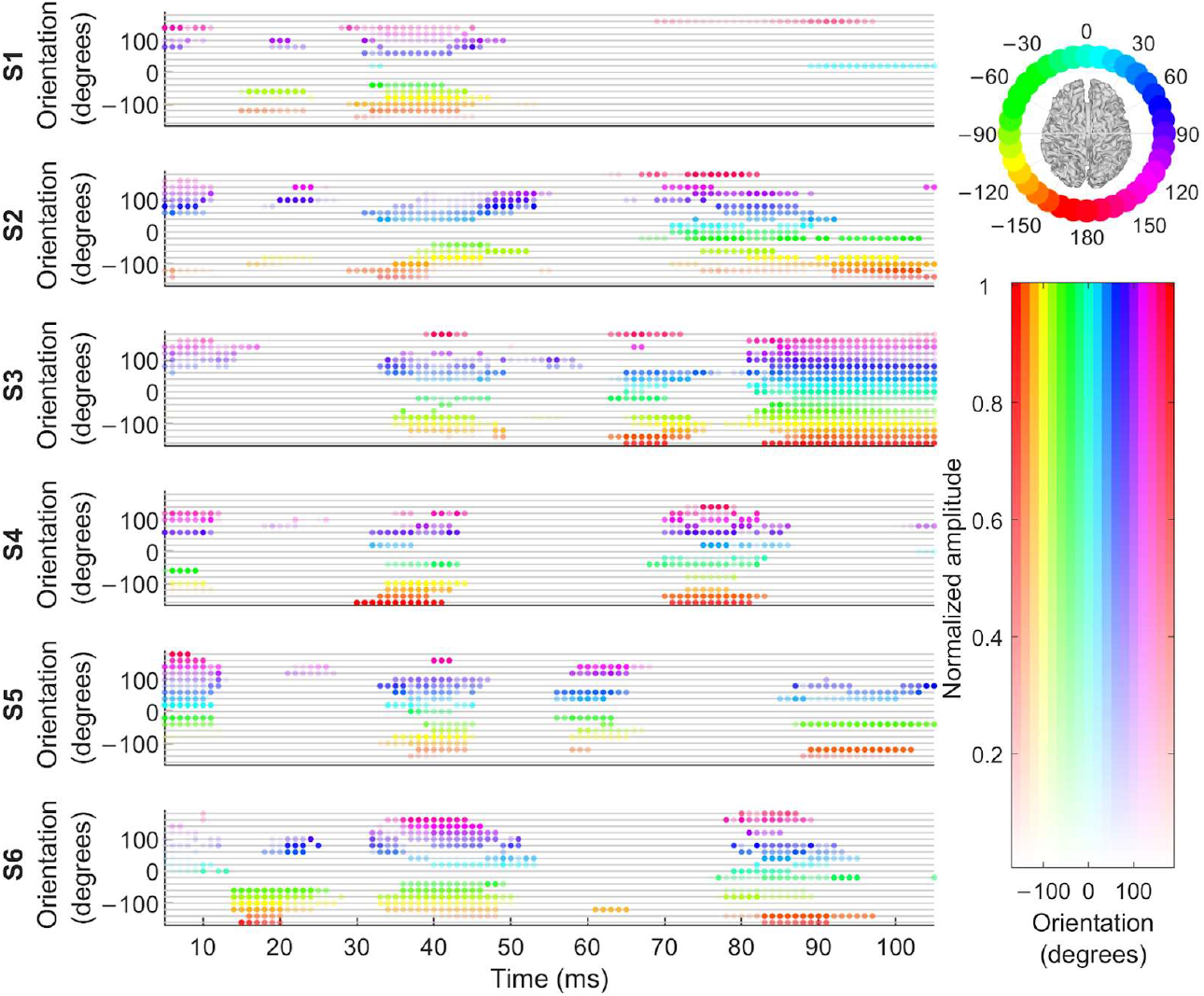
For each time point, the orientations producing a suprathreshold source are indicated as colored dots, where the color shows the stimulus orientation, and the opacity maps the normalized amplitude of a source with maximum concurrent activation at each time point. The orientations around –90 and 90 degrees are more likely to produce suprathreshold sources in the time interval 15…50 ms in all subjects.

## 5 Discussion

We studied the effect of TMS electric field orientation on TEP amplitudes, induced oscillations, and TMS-evoked signal propagation. The amplitudes of the positive peak at 20 ms (P20) and negative peak at 40 ms (N40) post-stimulus were especially sensitive to the stimulating electric-field orientation in the pre-SMA. The early, negative peak at 6 ms also appeared to reflect stimulus orientation; however, this peak could potentially be attributed to TMS-elicited muscle artefacts [7].

The cluster-based statistical analysis suggested an effect of stimulus orientation on TEPs only for the first 100 ms after the TMS pulse, except for S3, and was most pronounced at the stimulation site. The most prominent TEP amplitudes were evoked when orienting the electric field perpendicularly to the gyrus, in line with previous research [13,15]. In contrast to corticospinal responses [9], the visual inspection of the TEP amplitudes did not reflect prominent changes with a 180° shift in stimulus orientation, except for the potentially artefactual 6-ms peak. This might be attributed to EEG measuring postsynaptic currents, which are more generic to all cortical activations compared to corticospinal activations. In other words, due to the spatial limitation of EEG, 180° shifts in the orientation might produce similar responses despite potentially activating slightly different neuronal populations. This is further supported by the fact that the topographies of the responses (see supplementary material) do not change much with orientation. Thus, different stimulus orientations either activate similar neuronal populations and only affect the stimulation efficacy, and/or, EEG cannot distinguish between the post-synaptic currents from different neighbouring neuronal populations.

Although the sizes of the significant clusters do not necessarily exactly reflect the size of the effects studied, the analysis suggested that three subjects (S2, S3 and S6) showed some changes in TEPs in the contralateral hemisphere, and three subjects (S3, S5 and S6) showed changes in TEPs above the primary motor cortex at 50–100 ms. However, these changes were of smaller amplitude than those above the pre-SMA. Notably, only one subject showed indications of any effects of the stimulus orientation after 100 ms in the average TEP, indicating that the later responses may reflect multisensory responses or other non-specific responses to the TMS pulse, or may be more variable than earlier components. The time–frequency analysis supported the TEP findings of strongest changes in beta and gamma bandpower in the channels around the stimulated site right after the TMS pulse. This is consistent with the findings of Rosanova et al. [35], who reported evoking fast beta-gamma band oscillations when stimulating the superior frontal gyrus.

The methods for combining data from multiple subjects gave consistent results with the subject-level analysis and could potentially be utilized in future EEG analyses with several subjects. Fisher’s and Stouffer’s Z methods may offer a way to conduct population-wise analyses with less averaging over individual differences in comparison to directly combining data across participants.

The Activation Concurrence appeared highest at 35–45 ms in all subjects, while displaying large changes in the response amplitudes. A few subjects also showed more reliable responses around the stimulation site at 5–15 ms (S2 and S5) and 25–35 ms (S1, S4 and S6). Notably, the inter-condition variability appeared lower in this time interval as compared to 35–45 ms. After 55 ms, for all subjects except S1 and S3, only certain orientations evoked suprathreshold responses in central areas (S2, S4 and S6) and over the motor cortex (S5). In S3, almost all orientations evoked widespread suprathreshold activity at 75–105 ms, but this suprathreshold response spread to the contralateral frontal areas only when stimulating in certain orientations. These results indicate that the orientation may also play some role in efficacy of signal propagation from the stimulated site, in addition to influencing the efficacy of stimulation in terms of response amplitudes. If true, the coil orientation might after all influence which neurons are primarily stimulated by the induced electric field as suggested by [18,19], therefore playing some role in selecting which networks are engaged by the stimulation.

## 6 Conclusion

The stimulus orientation has a large effect on the P20–N40 peak-to-peak amplitude when stimulating pre-SMA. Analysis of variance (ANOVA) suggests orientation dependency in the data until approximately 100 ms after the TMS pulse, and changes in the power in the beta and theta bands. Source estimation suggested changes in both response amplitudes and signal propagation. These results converge with and expand on previous research that explored the early peaks after stimulating in different orientations. The methods applied here are promising for exploring the effects of multiple conditions with TMS–EEG data.

## Appendix A

### Fisher’s and Stouffer’s methods

Fisher’s method assumes that several weak observations can be combined into one stronger one. The combined test statistic can be calculated as follows:

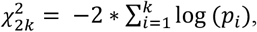

where *pi* indicates the *p*-value of the individual tests and *k* is the number of tests. Under the null hypothesis, the test statistic follows a Chi-squared distribution with 2*k* degrees of freedom, the null hypothesis being that all tests are true under their respective null hypotheses. The alternative hypothesis is that at least one of the null hypotheses is false.

Stouffer’s Z method [28] relies on combining the Z scores of the individual test into one test statistic:

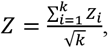

where *Z*_i_ are the Z scores of the individual tests, and *k* is the number of tests.

Fisher’s method was more sensitive than Stouffer’s Z method, as effects in single subjects can be sufficient to reach overall significance with the method. Stouffer’s Z method, on the other hand, is more sensitive to weaker effects present in several subjects.

#### Cluster-based statistical tests with Fisher’s and Stouffer’s methods

For applying Fisher’s and Stouffer’s Z method in the cluster-based permutational framework, first, at each permutation in, the trials from different orientations were shuffled within each subject to form a random partition containing all 18 orientations. Next, for each partition, we calculated the ANOVA scores at each channel time point within each subject, with the orientation as the independent variable. Finally, Fisher’s or Stouffer’s methods were applied at each channel time point to combine the subject-specific ANOVA statistics into a group-level measure. The final clustering was done on the combined statistics.

#### Source estimation

To calculate the lead fields that describe the sensitivity profiles of the EEG channels to cortical sources, the scalp, skull and white-matter surfaces were extracted from the subject-specific MRIs with the *headreco* [36–38] function of the SimNIBS software [33]. The surface meshes were decimated to ∼20,000 nodes and cleaned from surface artifacts with the iso2mesh package [39]. The boundary element method was utilized for calculating the lead-field matrices, assuming conductivity values of 0.33, 0.0033, and 0.33 S/m for the scalp, skull, and intracranial cavity, respectively. The 1/100 ratio between tissue types compensates for disaccounting the cerebrospinal fluid [40]. For calculation of the MNE estimates, the regularization parameter was set for each individual with Morozov’s discrepancy principle, by finding the regularization parameter that satisfies

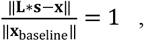

where **L** is the lead field matrix, **s** is the regularized source estimate, **x** the original sensor data from 0 to 300 ms averaged across trials, and **x**baseline the data from –300 to 0 ms averaged across trials. The found regularization parameters ranged from 0.08 to 0.3. The minimum-norm estimates were depth-weighted [33] with a depth parameter of 0.3, and the dipoles were placed with a free orientation in the lead field matrix.

### Declaration of interest

No authors have competing interests to declare.

### Funding sources

This project has received funding from the European Research Council (ERC) under the European Union’s Horizon 2020 research and innovation programme (grant agreement No. 810377) and from the Wellcome Leap as part of the Multi-Channel Psych Program. Additionally, funding has been received from the Swedish Cultural Foundation, and the Jane and Aatos Erkko Foundation. VHS received funding from the Research Council of Finland (decision No. 349985).

