## Supplementary material for "The dependency of TMS-evoked potentials on electric-field orientation in the cortex"

The visual inspection of the EEG data at different channels and time points confirmed that the assumptions of single-factor ANOVA hold for the data. Figure 2 illustrates a small sample from S1.

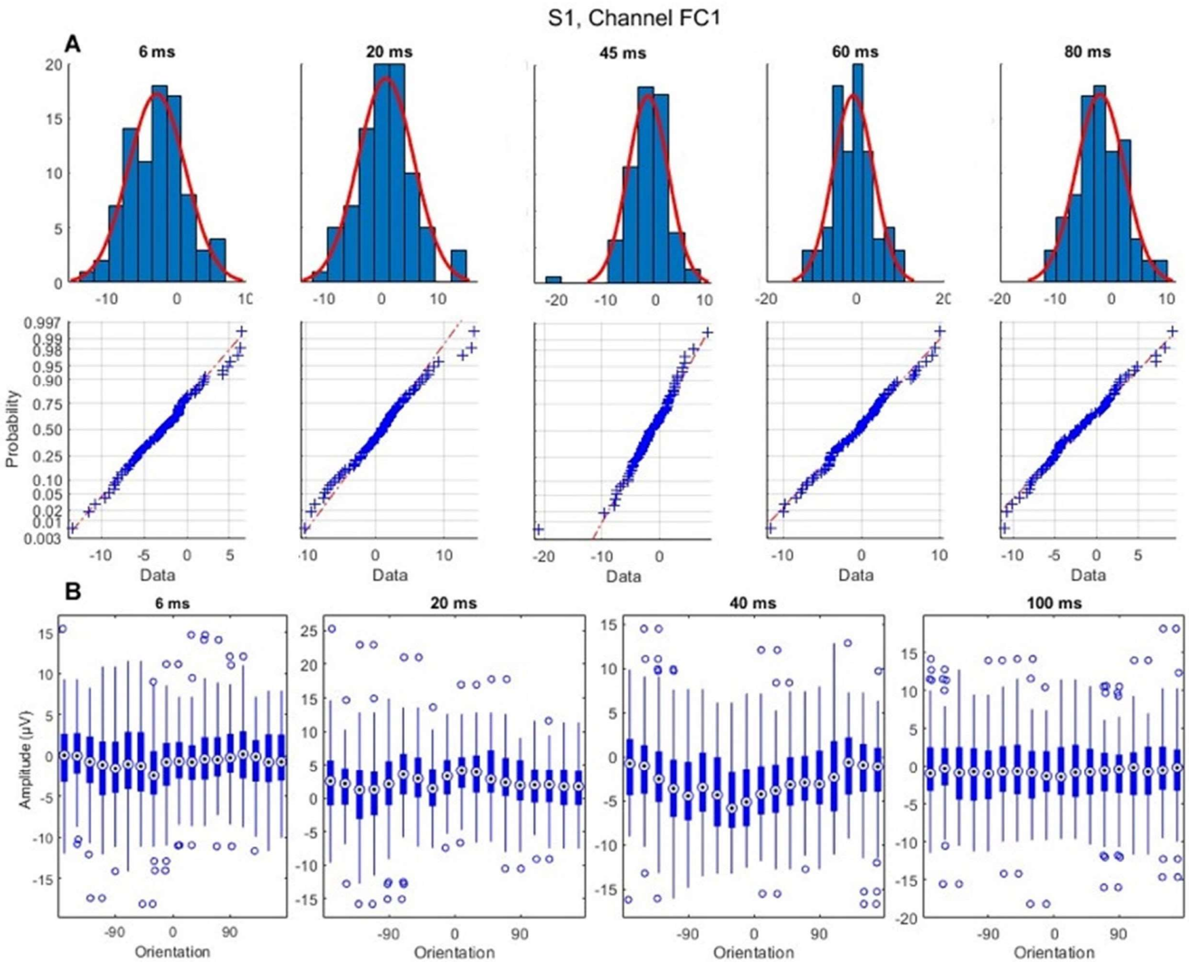


**Figure S1. A**. Histograms with fitted normal distributions and respective normal probability plots of the EEG data (S1, FC1) across trials of one orientation. The data appear close to normal, with some mildly heavier tails. **B**. Boxplots of the EEG data across trials in different orientations. The variances across orientations vary slightly, but are not visually different.

Topographies of averaged responses from –130°, –90°, –40°, 0°, 40°, 90°, 130° and 180° (±10°) were visualized for time points 20, 40, 60, 80, 100 and 150 ms. Visual inspection of topographies across different orientations suggested that the shapes of the early responses do not change much when changing the orientation. Especially at time points of large amplitudes, changes in the topographical distributions are small. On the other hand, the amplitude of the responses depend strongly on the orientation.

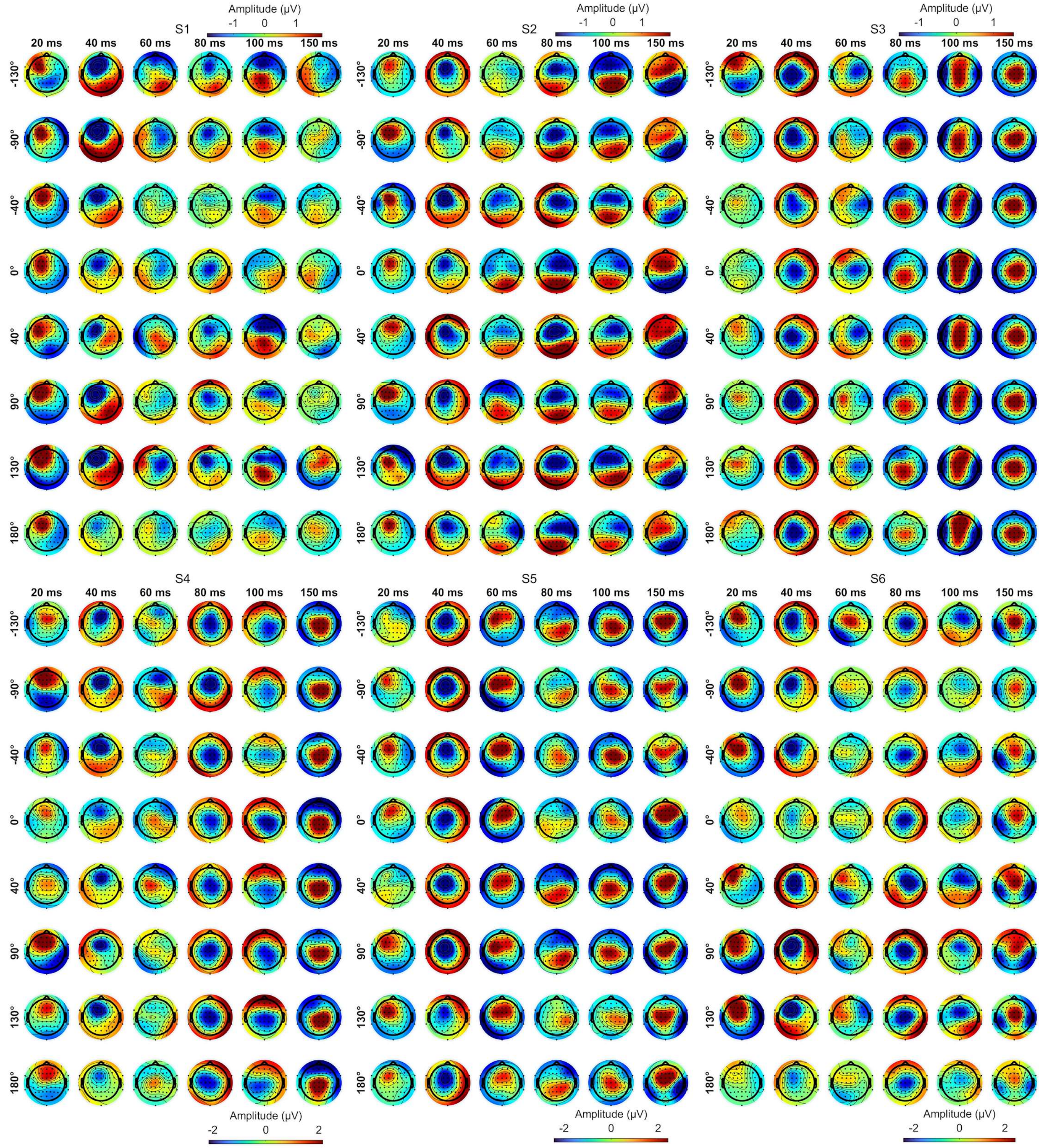


**Figure S2.** Visually inspecting responses from various orientations confirm that for the most part, the orientation affects the topographies mostly to a smaller degree, especially when response amplitudes are strong.
